# Phage biocontrol reduces the burden on plant immunity through suppression of bacterial virulence

**DOI:** 10.1101/2025.04.26.650785

**Authors:** Sebastian H. Erdrich, Milan Župunski, Ulrich Schurr, Guido Grossmann, Julia Frunzke, Borjana Arsova

## Abstract

Bacteriophages are increasingly recognized as key players in modulating plant-microbe interactions, including their potential in the biocontrol of plant pathogenic bacteria. In this study, we investigated the tripartite interaction between *Arabidopsis thaliana*, the bacterial plant pathogen *Xanthomonas campestris* pv. *campestris* (*Xcc*), and the lytic phage Seregon. Using parallel transcriptomic profiling, we characterized host and pathogen responses during infection and phage treatment. While treatment with phage Seregon did not lead to the eradication of *Xcc*, it significantly mitigated *Xcc*-induced disease symptoms, restoring leaf area to levels comparable to the uninfected control within 14 days post-inoculation. Our data revealed that phage-mediated protection is associated with early bacterial recognition, and suppression of Jasmonate (JA)-related responses in the host. Analysis of nuclear localized reporter plant lines further confirmed a significant reduction in ROS levels and JA biosynthesis in phage-treated plants. Concurrently, *Xcc* exhibited significant transcriptional downregulation of key virulence factors in the presence of the phage, including the genes encoding the type III secretion system, its associated effectors, and components involved in flagella biosynthesis. Remarkably, phage treatment did not lead to a significant increase in bacterial resistance to phage infection, which is in stark contrast to *in vitro* conditions. Taken together, this study provides first mechanistic insight into how phages can be harnessed to shape plant-pathogen interactions and highlights their potential role in enhancing plant resilience through targeted modulation of both host immunity and pathogen behavior.

## 1. Introduction

The escalation of bacterial plant pathogens [1], coupled with the diminishing effectiveness of traditional antibacterial treatments due to overuse and shifting climatic conditions, threatens our food supply [2, 3]. The increasing prevalence of antibiotic-resistant pathogens adds to the concern. In this context, microbe-based strategies present sustainable and effective alternatives to antibiotics, chemical fertilizers, and pesticides, which can have considerable impacts on human health and environmental sustainability.

Bacteriophages (phages) are specialized viruses that infect and eliminate bacteria. Their therapeutic potential in treating human or plant diseases was recognized early after their discovery more than a century ago [4]. However, interest declined with the rise of highly effective chemical biocides that offered broad-spectrum applicability. In the light of the growing antibiotic resistance crisis and the impacts of climate change, phage-based solutions are now regaining significant attention for their precision, effectiveness against plant pathogens, and safety for agricultural applications [5, 6]. The advancement of these environmentally sustainable alternatives is reinforced by policies like the European Union’s Green Deal, which targets a 50% reduction in synthetic agrochemical use by 2030 (https://food.ec.europa.eu/plants/pesticides/sustainable-use-pesticides_en). Versatile applicability of phages is shown by recent studies reporting on the development of effective phage-based sprays, soil treatment formulation and seed coatings [7]. Efficient phage-based biocontrol was demonstrated for some of the major plant pathogens, including *Pseudomonas syringae, Xanthomonas campestris, Xanthomonas oryzae, Erwinia amylovora, Ralstonia solanacearum, Pectobacterium carotovorum* and *Xylella fastidiosa* [6, 8].

Constantly exposed to various pathogens of bacterial, fungal or viral origin, plants have developed intricate sensing and regulatory networks to fine-tune their immune response. The classic ZIG-ZAG model of plant immunity describes a multi-layered defense system featuring sequential activation to protect the plant [9]. If one layer fails, the activation of a subsequent layer can still provide resistance or, if ineffective, lead to susceptibility. The first layer of defense detects microbe-associated molecular patterns (MAMPs) via pattern recognition receptors (PRR), which recognize conserved bacterial features like cell wall components [10–12] or flagellin [13–16]. One prominent example is the detection of the Flg22 region, though some *Xanthomonas* strains evade this recognition by modifying Flg22 [17]. Detection by PRRs gets transduced by a cascade of mitogen-activated kinases (MAP-kinases)in the second layer of defense [18–20], finally leading to transcriptional activation of defense genes and the so-called microbial-pattern-triggered immunity (PTI). Pathogenic bacteria, including *X. campestris* (*Xcc*), deploy type 3 secretion systems (T3SS) to deliver effector protein into the plant cell [21–25] to interfere with the PTI signalling cascade [26]. This defense layer is tuned to detect these effectors, either directly or indirectly [27, 28], and to activate heavy defense responses, e.g. production of reactive oxygen species or induction of cell death to prevent further spreading of the disease. If successful, this leads to effector-triggered-immunity (ETI) [29]. ETI also plays a crucial role in the interaction between *Xcc* and *Arabidopsis* [30].

Local immune activation translates into systemic responses through defense-related plant hormones, including salicylic acid (SA)[31], jasmonate (JA)[32, 33], ethylene (ET)[34] and abscisic acid (ABA)[35]. These hormones induce systemic acquired resistance (SAR) by activating defense genes beyond the infection site [36]. However, defense responses are energy-intensive, creating a trade-off with growth. Plants have one last trick up their sleeve: close monitoring of their environment and detection of bacterial quorum sensing molecules like N-acetyl-homoserine lactones, leading to low-level defense activation across the whole plant, enabling a rapid response upon pathogen attack [37].

In this study, we focused on the biocontrol of *X. campestris (Xcc)*, a major plant pathogen responsible for black rot disease in *Brassicaceae* [38]. *Xcc* is a seed-transmitted bacterium with both epiphytic and endophytic vascular life phases. Under favorable conditions and at high population densities, it enters the plant through natural openings such as hydathodes, stomates, cracks or wounds [39, 40]. This collective invasion is coordinated by diffusible molecules and recognition systems, involving chemoattraction and quorum sensing systems [41]. Once inside the plant, *Xcc* establishes biofilms in the xylem, drawing nutrients from the sap. By clogging vascular bundles with biofilms and short-circuiting plant defense responses with effector proteins, *Xcc* causes the typical V-shaped necrotic lesions in important crop plants like *Brassica oleracea*.

While numerous historical and recent studies highlight the potential of phage-based biocontrol, the impact of phage treatment on plant immune responses and bacterial adaptation to phage predation within the tripartite interaction remains entirely unexplored. In this study, we established a gnotobiotic system to study the impact of phage biocontrol on *Arabidopsis thaliana* infected with *Xcc*. Here, a single treatment with our previously described lytic phage Seregon [42] infecting *Xcc* resulted in the restoration of healthy plant growth in a period of two weeks post infection. Parallel transcriptomic profiling at different time points after infection revealed a significant plant immune response to *Xcc*, which was reduced under biocontrol conditions. While bacteria were not completely eradicated by a single phage treatment, expression of bacterial virulence and motility genes was significantly reduced. Remarkably, treatment conditions did not lead to the emergence of high levels of bacterial resistance to phage Seregon. These data offer first molecular insights into the complex interplay between plant, bacteria and phages, advancing research to the next level of understanding these intricate interactions.

## 2. Material and Methods

### 2.1 Plant propagation

*Arabidopsis thaliana* Col-0 plants were propagated on ½ Murashige and Skoog-Agar [43]. Individual surface sterilized seeds were placed in a petri dish in a 3×3 matrix under a laminar flow bench. Plates were sealed with micropore tape and put into a climate chamber with a 12 h light–12 h dark regime at 22°C day and 18°C night, with 100 μE illumination at a relative humidity of 60%.

### 2.2 Bacterial growth conditions

*Xanthomonas campestris pv. campestris* str. 8004[38] was used in this study. Cultures were grown on plates and inoculated from single colonies in liquid media for overnight cultures. Bacteria were grown on nutrient agar (5.0 g peptone, 3.0 g yeast extract and 15.0 g Agar in 1,000 ml of dH_2_O). Unless mentioned otherwise, bacterial cultivation was performed in shake flasks at 28°C and 150 rpm.

### 2.3 Phage amplification

The *Xanthomonas* phage Seregon was isolated and described previously [42]. Phages were either amplified on plates or in liquid cultures. For plate amplification, a double agar overlay was prepared using 100 µL phage solution with a titer 10^8^ Pfu/mL added to 3.5 mL 0.4%-top agar containing the bacterium at a final OD_600_ of 0.2. Harvesting of purified phage particles was performed after overnight incubation. The top agar was solubilized by adding 3 mL ½ MS-Buffer followed by two hours of incubation on a rock shaker. The solution was subsequently transferred into a falcon tube and centrifuged at 5000 *g* for 25 min to remove residual amounts of agar and bacterial cell fragments. The supernatant was filtered twice through 0.2 µM syringe filters and stored at 4°C. For titer determination, a dilution series was spotted on overlay agar, and the visible plaques at the highest dilution were counted to determine the plaque-forming units (pfu) per mL.

### 2.4 Plant inoculation

Inoculation of plants was performed by flood inoculation, similar to Korniienko et al. 2021 [44]. Eight-day-old seedlings were submerged within 50 mL ½ MS liquid medium containing 10^9^ cfu/mL *Xcc* for the pathogen condition. Both phage and bacterium were purified via centrifugation and resuspended in ½ MS, a small volume of this higher concentrated stock was used to adjust the desired concentration for the inoculation solution. For the phage treatment condition, phages were added to the solution before flooding, at a multiplicity of infection of 5 (MOI5 = five phages per bacterial cell). The phage-only control was flooded with 50 mL ½ MS containing the same amount of phages as in the MOI 5 treatment. The plant-only control was flooded with ½ MS. After 5 minutes, the liquid was removed carefully, and the plates were again sealed with micropore tape and incubated in the climate chamber. Plant growth in the different treatment conditions was consistent over 5 independent experimental rounds; and all biological samples used for further analysis were generated in triplicates.

### 2.5 Non-invasive plant phenotyping

The non-invasive plant phenotyping was performed at 0, 2, 5, 7, 9 and 14 days post-infection (dpi). Plates were placed onto an Epson 10000XL WinRhizo scanner, and images were acquired using the manufacturer’s scanner interface. The top of the plates was covered with white cardboard for better contrast. Tiff images were retrieved by scanning from below, with a resolution of 1200 dpi. The plant leaf area was calculated using Image J (Version 1.54f), and green pixels were isolated by HSV thresholding. The leaf area of each plant was determined using the “analyse particle” function.

### 2.6 Invasive plant phenotyping and extraction of microbes

The invasive plant phenotyping was performed in a separate experiment and plants were harvested at 0, 2, 5, 7 and 14 (dpi). The fresh weight of plants was determined by placing plants into pre-weighed microcentrifuge tubes containing one 3 mm glass beads using sterile tweezers. For the duration of the experiment, samples were kept on ice. The fresh weight was determined by subtracting the pre-weighed weight of the tube including the glass bead-weight from the final weight. Afterwards 1 mL of ½ MS media was added and the tissue disrupted by using a ball mill for 10s at 60Hz/s. Afterwards, a dilution series was prepared for each sample. To determine the bacterial load 3 µl of each dilution step were plated on NB-Agar plates and CFUs were determined after 48 h incubation at 28°C. To determine the amount of infectious phage particles (pfu, plaque forming units), a double agar overlay was prepared as described above and 3µl µl of each dilution step were plated on top. Phage titer was determined after 24 h incubation at 28°C.

### 2.7 RNA extraction

For RNA extraction, multiple plants with a total weight of 45 mg were harvested from each treatment group and pooled in a microcentrifuge tube. The obtained tissue was immediately frozen in liquid nitrogen and stored at −80 °C. Sampling occurred in triplicates at 0, 2 and 7 dpi. For nucleic acid extraction, the frozen tissue was ground in a ball mill (Retsch MM200, Retsch, Germany) with a 4-5 mm stainless steel beat. Total RNA extraction from the resulting tissue powder was performed using the Rneasy^®^ Plant Mini Kit (Qiagen, Hilden, Germany) according to the manufacturer’s instructions. To remove genomic DNA contamination, total RNA was incubated with DNAse I (Thermo Fisher Scientific, AM1906) for 15 min.

### 2.8 RNA-Seq

At the start of the experiment, each condition contained 216 plants; the plants for RNA extraction were harvested throughout the experiment at 0, 2, and 7 days post-inoculation. RNA-Seq was performed for samples of the conditions: control, phage only, *Xcc* and MOI5 at 2 and 7 dpi, each in three biological replicates as well as one triplicate of plants directly before treatment (0dpi) was performed with rRNA depletion strand-specific RNA library preparation at Azenta Life Science, followed by RNA-Seq on a Novaseq 6,000 with a sequencing depth of 100M read pairs per sample (2x 150 bp). Raw read sequences were archived at NCBI (GSE294853). The read quality was assessed with the FastQC software v.0.11.9 (http://www.bioinformatics.babraham.ac.uk/projects/fastqc/). Sequencing adapters and low-quality sequences were removed using the Trimmomatic tool v. 0.39 [45]. The obtained reads were mapped against the *Arabidopsis thaliana* genome (TAIR 10.0)[46] using Kallisto [47]. Genome mapping against the other species (*Xcc* [NC_007086.1] and phage Seregon [ON189048.1]) was performed in subsequent rounds using Kallisto. Differential expression analysis was performed using DESeg2 [48]. The DESeq2 Algorithm attained the differentially expressed genes using the Wald test and DEGs were corrected for multiple testing using the Benjamini and Hochberg method. The significance cut off were set to <0.05 p-adj and LogFC > 1.

### 2.9 Imaging of nuclear reporter lines

For microscopic analysis of pAOS::NLS-3xVenus and pPER5::NLS-3xVenus reporter lines [49] seven-day-old seedlings were carefully transferred onto ½ MS agar medium flood inoculated as described above with *Xcc*, phage Seregon, or MOI5 solution, and grown for 48 hours prior to imaging. The seedlings were observed under spinning-disk confocal microscope 48 hours post inoculation.

Imaging of plant roots was done using a custom-build fluorescent microscope with a Nikon Ti-E stand, equipped with a Nikon 10X objective (S Fluor, N.A.=0.50), CrestOptics X-light spinning disk (pinhole size 50µm), motorized stage (Applied Scientific Instrumentation, USA), motorized filter wheel (Cairn Research, UK), laser launch (Omicron, Germany), dichroic mirror (Chroma triple band 440/514/561), and an sCMOS camera (Teledyne Photometrics, Prime 95B, USA). Image acquisition was operated through Nikon NIS Elements 4.0 software. Imaging of reporter lines and PI staining were done with 520nm excitation (80mW power), with a 542/20 emission filter for reporter lines, and 605/70 emission filter for PI staining. After acquisition, the data was processed and analyzed with ImageJ/Fiji (version 1.49b), following the “Subtract background” (rolling ball radius = 200px) function, and the “Gaussian Blur” (sigma = 1 radius) filter. Maximum intensity projections were generated for each dataset. At least two independent experiments were performed, with 6-12 replicates analyzed. Sample numbers AOS: Mock 13; Seregon 13; Xcc 22; MOI5 26. PER5: Mock, Seregon, Xcc, MOI5 = 7. Data were analyzed with non-parametric test, due to non-normal distribution and unequal variances. Overall group differences were assessed using the Kruskal–Wallis rank-sum test (stats package), followed by pairwise comparisons with Dunn’s test (FSA package, v0.9.4) and Benjamini–Hochberg correction. Significance groupings were visualized using compact letter displays from the multcompView package (v0.1-9).

## 3. Results

### 3.1 Phage biocontrol restores plant growth in the presence of the bacterial pathogen *Xanthomonas campestris*

To investigate phage-based biocontrol in a defined laboratory setting, we established a tripartite system consisting of the model plant *Arabidopsis thaliana*, the bacterial plant pathogenic organism *Xanthomonas campestris pv. campestris* (*Xcc*) and phage Seregon infecting *Xcc* [4]. For infection experiments we used a plate-based gnotobiotic system (Figure 1 A; B). While infection of *A. thaliana* with *Xcc* resulted in a significant reduction in leaf area and plant fresh weight, treatment with the phage strongly reduced the adversary effect of *Xcc* on *A. thaliana*. First phenotypic differences were noticed after 5 dpi. At 12 dpi, a significant difference in the leaf area and plant fresh weight was observed between the control and the *Xcc* treatment (Figure 1 C; D). Throughout the experiment (0-14 dpi), there was no significant difference between the leaf area of the control, the phage-only condition and the phage biocontrol treatment against *Xcc* at a multiplicity of infection of 5 (MOI5 = five phages per bacterial cell).

**Figure 1.**
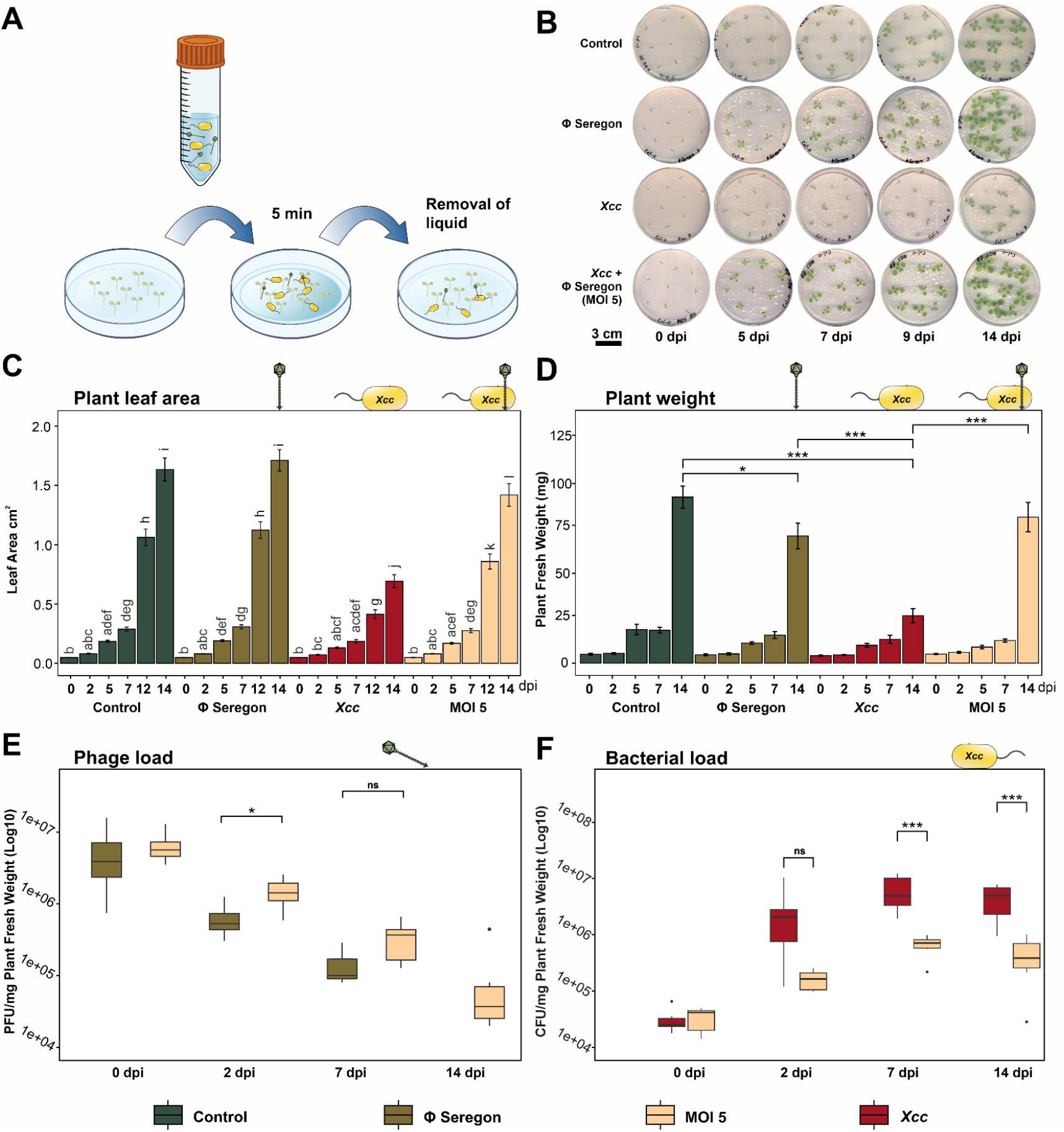
Phage mediated plant protection over time - Tripartite interaction in a gnotobiotic system and plant phenotyping. **A)** Inoculation workflow. **B)** Depiction of the plants over time showing the control-inoculated plants (first row, ½ MS media only), *X. campestris* (second row), *Xcc* plus phage at MOI5 (third row), phage Seregon (forth row) **C)** Non-invasive measurement of leaf area over time. ANOVA test followed by Tukey HSD F(3.234)=31.25, p=0.0001. **D)** Fresh weight per plant over time. **E)** Phage load per mg of plant fresh weight. **F)** Bacterial load per mg of plant fresh weight. (N=6 plants for 0, 2, 7 dpi and N=10 for 14 dpi).

To quantify the effect of phage biocontrol on the abundance of bacterial cells and infectious phage particles, we determined bacterial cfu/mg plant fresh weight as well as phage Seregon pfu/mg (Figure 1 E). We observed a ten-fold reduction of bacteria over the course of the experiment. Phage titers declined over the course of the experiment, but stayed consistently higher in the biocontrol condition (presence of *Xcc* as host) in comparison to the phage-only control. While the bacteria were not eradicated by this single phage treatment, plant growth was nevertheless restored to the level of the uninfected control underlining the efficacy of phage pressure in this setting and its potential for biocontrol applications.

### 3.2 Tripartite transcriptomic analysis of phage-based biocontrol

To gain insights into plant defense signaling and bacterial response to phage treatment, we performed a comprehensive tripartite parallel transcriptomic study. All samples were processed via a double rRNA depletion for plants and prokaryotes, prior to sequencing. Across all conditions, we detected 76.25% of all *Arabidopsis* genes, 72.74% of *Xcc* genes and 91.69% of phage Seregon genes. Only in one sample, at 7 dpi, no phage reads were detected.

PCA plots of plant samples show clear separation of treatment and time-point dependent changes (Figure 2A). To understand how *Arabidopsis* responds to *Xcc* in the presence or absence of phage Seregon, we performed an analysis of differential expressed genes (DEGs) of each treatment compared to untreated plants at the same developmental time point. In line with the phenotypic data (Figure 1 B) pathway enrichment analysis using the Kyoto Encyclopedia of Genes and Genomes (KEGG) database revealed 17 immune-related pathways upregulated in *Xcc-*treated plants at 2 dpi (Figure 2 C upper panel). In contrast, phage treated samples only revealed 3 upregulated pathways. By 7 dpi, seven pathways remained enriched, with a role in secondary metabolite production. To further visualize the effect of phage biocontrol on plant responses we compared DEGs and looked for enriched pathways during biocontrol. Pathway enrichment showed that 3 distinct pathways are downregulated under biocontrol conditions compared to *Xcc* infected plants at 2 dpi (“carotene catabolic process”, “terpene catabolic process” and “isoprenoid catabolic process”), whereas at 7 dpi only one pathway was upregulated (“RNA modification”, Figure 2 C). The carotenoid pathway is linked to ABA biosynthesis and redox regulation [50]. Terpenes, which are plant secondary metabolites often induced by elicitors, exhibit antimicrobial activity [51, 52], similar to isoprenoids – particularly those derived from the mevalonate (MVA) pathway [53]. The observed downregulation of these 3 pathways in phage-treated plants suggests a suppression of immune responses compare to plants infected with *Xcc*. Notably, some of the most strongly downregulated genes in phage-treated, infected plants include key regulators of programmed cell death (BAP2), ABA signaling (CML 37), redox regulation (AOX1D) and immunity regulation via JA (WRKY59) (Figure 2 D).

**Figure 2.**
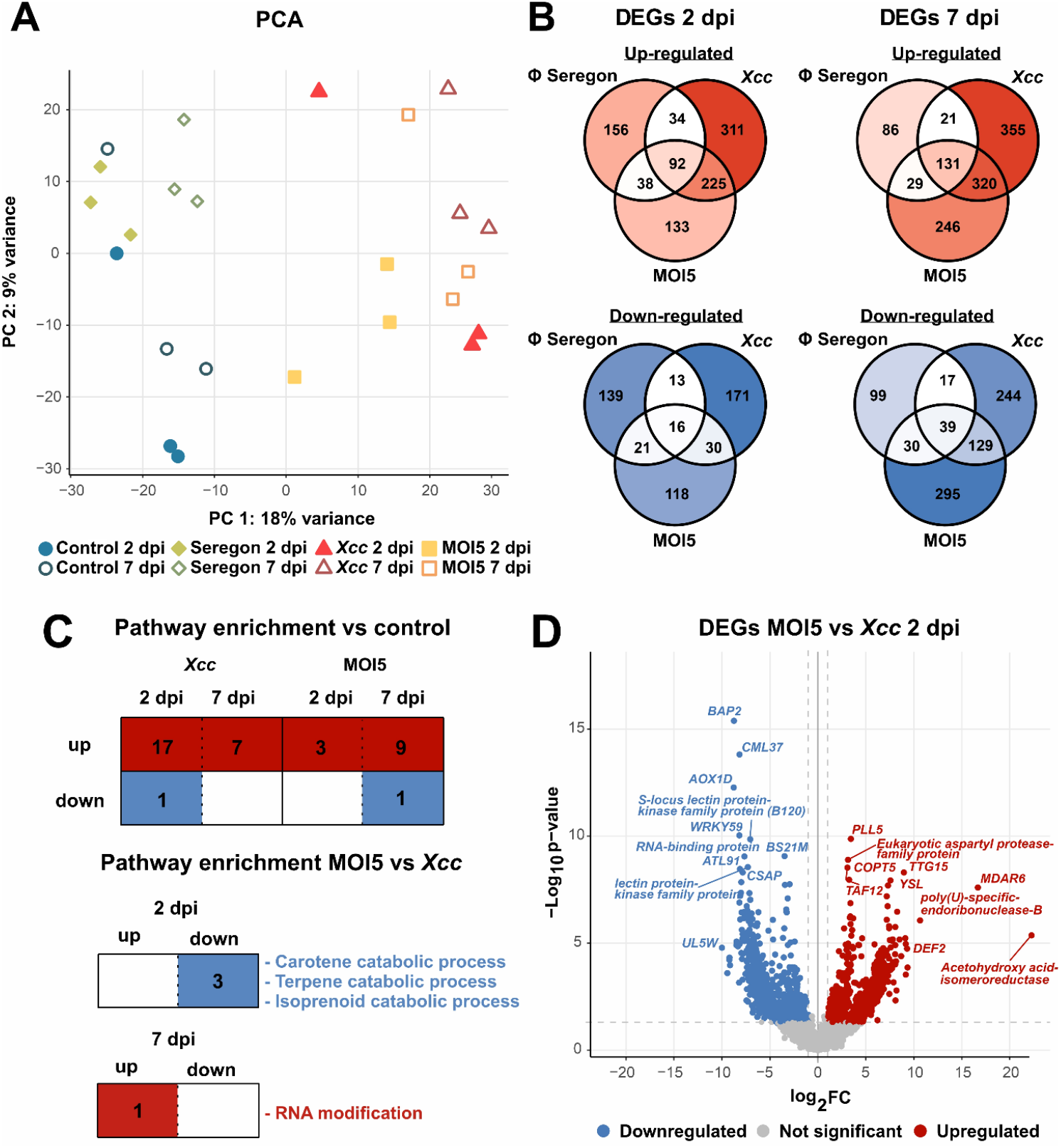
Differentially expressed *Arabidopsis* genes (DEGs) point towards a reduced need for full defence activation during tripartite interaction. Significant differentially expressed genes were identified using the DESeq2 algorithm [48]. **A)** Principal component analysis of expressed plant genes. Samples are coloured according to their treatment condition. **B)** Venn diagram of differentially expressed genes during tripartite interaction. Upregulated genes are displayed in red, downregulated genes are displayed in blue. Each condition was tested against the control at the respective development time point which was used as a reference (p-value <0.05). The resulting condition-specific differentially expressed genes were then compared to every other condition to see if there is an overlap. **C)** Significantly enriched KEGG pathways (Supplemental Table S1). Upper table: compared to day-specific control; Lower table: biocontrol compared to *Xcc* infected plants. **D)** Volcano plot of phage treatment (MOI5) compared to *Xcc* conditions, 2 days post inoculation. Downregulated genes are displayed in blue, whereas upregulated genes are shown in red. Dotted grey lines indicate the significance threshold (p-value <0.05 and Log2 foldchange > 1). The ten most significant differentially expressed are displayed with gene names.

### 3.3 Infection with *Xcc* triggers significant plant defense signalling

Upon *Xcc* infection, *Arabidopsis* exhibited a strong immune response reflected on the level of transcription (Fig 2B/C). Both pattern-triggered immunity (PTI) and effector-triggered immunity (ETI) are strongly upregulated at 2 and 7 dpi, signaling active pathogen recognition and defense activation (Figure S1). Notably, a broad range of nucleotide-binding leucine-rich repeat (NLR) proteins are induced, with *TIR-NBS6* (AT1G72890) showing a Log2FC ∼7.6 at 2 dpi, highlighting a robust ETI response.

During signal integration, key defense-related kinases are strongly upregulated. Several MAP kinases, cysteine-rich receptor-like kinases (CRKs), and transcription factors (WRKY, NAC, MYB) exhibited increased expression levels. *CRK13* and *CRK24*, implicated in cell death and defense signaling, showed strong induction at both time points. In particular, *CRK13*, which is linked to cell death initiation, was upregulated across multiple variants. Additionally, calcium signaling components, including *CAR1* (AT5G37740) and calmodulin *CAM8*, are significantly induced, supporting their role in amplifying the immune response.

Defense execution mechanisms are also activated, including genes involved in reactive oxygen species (ROS) production and lignification. Peroxidases such as *PRX2, IRR1*, and *PER28* exhibited increased expression, indicating elevated oxidative stress and cell wall fortification against *Xcc*. These findings underscore the extensive immune activation triggered by *Xcc*, characterized by strong PTI and ETI induction, ROS production, and transcriptional reprogramming toward defense.

### 3.4 Phage-based biocontrol leads to a reduction in plant immune activation

To gain deeper insights, which genes are particularly responsive to the presence of the phage-biocontrol (MOI5), we compared the transcript levels of phage-treated to *Xcc*-infected plants. Accounting for the p-value and the level of fold change, the 100 most significantly regulated genes are presented in Figure S2. The identified genes primarily fall into the following categories: PTI, ETI, oxidative burst, transcriptional activators, hormonal signaling, and cell death. The most striking regulation was seen in the expression of genes encoding a cutin synthetase (CUS2, AT5G33370), upregulated at 7dpi, a loricrin-like protein (AT5G09670) as well as MDAR6 (AT1G63940) involved in redox regulation upregulated at 2 dpi. Strongest downregulation was found in a cytosolic invertase (CINV1, At1g35580), involved in sucrose conversion to fructose and glucose.

A selection of transcripts, with functional annotations matching the three stages of plant immunity (immune recognition, signal transduction and defense execution pathways) as outlined in [54] is summarized in Figure 3 (numerical values in Supplemental Table 2). Among genes related to immune recognition, encompassing PTI, NLRs and ETI, we observed downregulation of PTI transcripts ATSBT3.3, PROPEP2- a MAMP receptor, LECRTK-IX.2 at both days, and of LECRTK-IX.2 at 7dpi. Interestingly BES1, which links bacterial response to the brassinosteroid signaling pathway [55] is upregulated 7 fold in MOI5 at both time points. In the group of NLRs, intracellular receptors that detect xenobiotic compounds within the cytosol, a number of Toll-Interleukin-Resistance (TIR) domain family proteins are downregulated in MOI5 plants. In the group of ETI transcripts, the genes SGT1A and At1g58420 are both downregulated in phage-treated plants at both time points.

**Figure 3.**
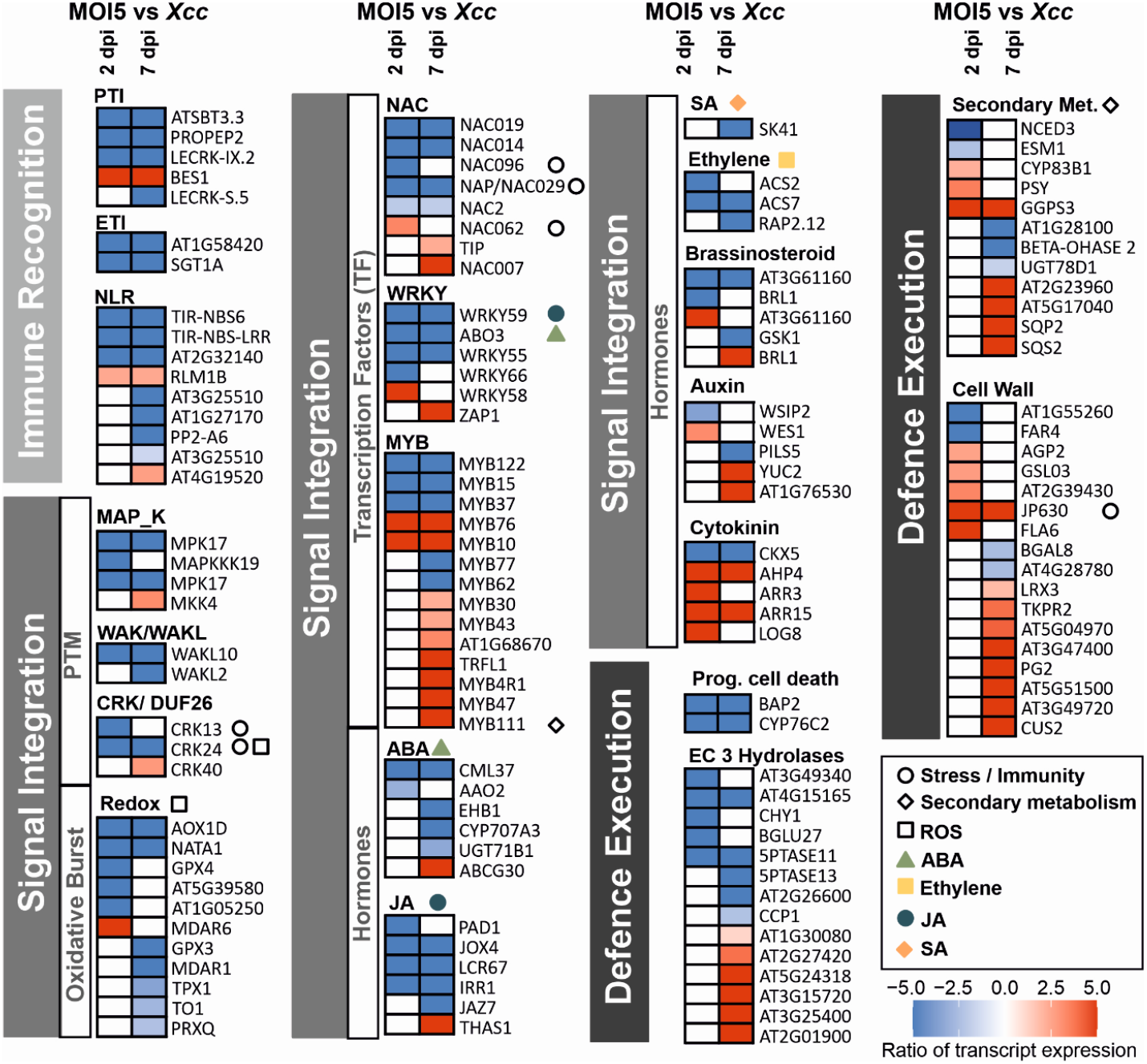
Phage biocontrol leads to altered defense regulation in *Arabidopsis* compared to the pathogen. Ratio of transcript expression of MOI 5 compared to Xcc infected plants (expressed as log2 (FC)). Shown are selected transcripts involved in pathogen recognition (light grey vertical bars) immune signal integration (medium grey) and defense execution (dark grey). The level of transcript expression is presented as a heat map, shown bottom right, with shades of blue indicating lower expression, and shares of red indicating higher expression. The heat map is cut off at +/-5 for visual simplification, numerical data can be found in supplemental table S2. Transcripts are grouped in functional groups based on MapMan mapping file X4.1 [66], with adaptations as listed in supplemental table 2. Where discussed in the text protein kinases, transcription factors, redox and hormone transcripts are linked through the use of symbols listed in the legend.

Signal transduction involves protein kinases (MAPKs, WAK/WAKL, CRK/DUF26), redox regulators, transcription factors (WRKY, NAC, MYB), and hormone signaling. Among the genes encoding transcription factors, we observed lower transcript levels for WRKY and NAC transcription factors (e.g., WRKY59, WRKY55, NAC019, NAC029), but increased expression of WRKY58, ZAP1, NAC062, and NAC007. NAC062 is relevant because it relocates to the nucleus under endoplasmatic reticulum stress and integrates biotic and abiotic stress responses, promoting cell survival [56] (Figure 3).

Notably, the group of MYB transcription factors deviates from the pattern of general downregulation during biocontrol. Here we see a constitutive downregulation of MYB 15, MYB37, and MYB 122. However, a much larger group of seven MYB transcription factors is upregulated at 7 dpiFigure 3. Among them, MYB111, is a master regulator of flavonoid synthesis [57], and could indicate shifts in secondary metabolism.

In the signal transduction subgroup related to hormonal signal integration, we selectively show the ABA, ethylene and jasmonic acid (JA) transcripts (full dataset-Table S6). The ABA signaling regulator CML37, a positive modulator of stress responses [58–60], is downregulated at both time points. Decreased ABA synthesis is indicated already at 2 dpi with a downregulated abscisic aldehyde oxidase (AAO2). Transcripts involved in ABA conjugation and degradation (abscisic acid UDP-glycosyltransferase UGT71B1, abscisic acid hydroxylase CYP707A3), as well as ABA perception and signaling (EHB1), are all downregulated at 7dpi. In contrast, ABA transporter ABCG30 is upregulated at 7dpi, Figure 3.

Among genes involved in auxin transport, *PILS5*, a growth inhibitor under stress conditions [61], is suppressed, while *PIN-LIKES4* is upregulated, potentially promoting normal growth (Supplemental Table 2). Ethylene signaling is downregulated constitutively in MOI 5 plants, with the ethylene synthesis related transcripts ACS2 and ACS 7 downregulated at 2 dpi and both time points, respectively. In addition, RAP2.12 involved in Ethylene signaling is downregulated at 7dpi.

Similarly, the jasmonate pathway (JA) showed a general downregulation. At 2 dpi, *PAD1*, required for jasmonate signaling [62], is downregulated in phage biocontrol compared to *Xcc* infected plants, along with IRR1, *JAZ7, Pdf1*.*1 and JOX4 –* a jasmonate induced oxygenase 4 (Figure 3). This is in line with the previously described downregulation of the jasmonate-responsive transcription factor *WRKY59*.

The final set of transcripts examined here belong to the defense execution step. Markers of hypersensitive response (HR) and cell death are notably downregulated, supporting our expectation that biocontrol limits excessive immune activation. *CYP76C2*, a cytochrome P450 involved in HR, is strongly suppressed (Log2FC -8.1). Similarly, *BAP2*, a key regulator of programmed cell death required for the IRE1-mediated UPR, is downregulated at 2 dpi (Log2FC -8.7 and -8.3*).

At the structural level, cuticle biosynthesis is reinforced through upregulation of *CUS2* [63] and a loricrin-like peptide [64], with lipid metabolism genes (*MBOAT, LPXA, ELT6, ALT1*) supporting suberization. RNA signaling is affected via *MORF1* upregulation, while the strong downregulation of *CINV1* suggests limited pathogen access to plant energy resources [65]. Additionally, programmed cell death regulators *CYP76C2* and *BAP2* are significantly suppressed.

Overall, these findings indicate that phage biocontrol reduces immune activation by lowering pathogen burden and preventing excessive defense-associated stress, allowing the plant to maintain growth and metabolic balance.

### 3.5 Nuclear localized reporter plant lines confirm reduced ROS and jasmonate response in phage-treated samples

One major driver of defense-associated stress is damage caused by free radicals during the generation of ROS. Transcriptomic data suggested reduced ROS production in the biocontrol treatment, with ROS inhibiting genes being downregulated after 2 dpi. For instance, the alternative oxidase 1D (AOX1D), which minimizes oxidative stress [67], is strongly downregulated (logFC -8.7), as is NATA1, a negative regulator of hydrogen peroxide. Additionally, cysteine-rich RLK (RECEPTOR-like protein kinase) 24 is significantly downregulated (log2FC -6.3).

To test the hypothesis that immune responses triggering ROS production are differentially regulated in presence or absence of phages, we used PER5:3xVenus-NLS lines of *Arabidopsis* [68]. Here the nuclear localized Venus reporter gene is fused to the PER5 promoter. PER5 encodes the microsomal ASCORBATE PEROXIDASE 5 involved in scavenging of hydrogen peroxide, resulting in increasing fluorescence signal in the nucleus under oxidative stress (Figure 4A/B). In our tripartite setting, *Xcc*-treated plants showed strong nuclear fluorescence accumulation in roots, while MOI5-treated plants exhibited a much weaker signal, consistent with lower ROS levels and corroborating the hypothesis of a reduced immune response under phage biocontrol conditions.

**Figure 4.**
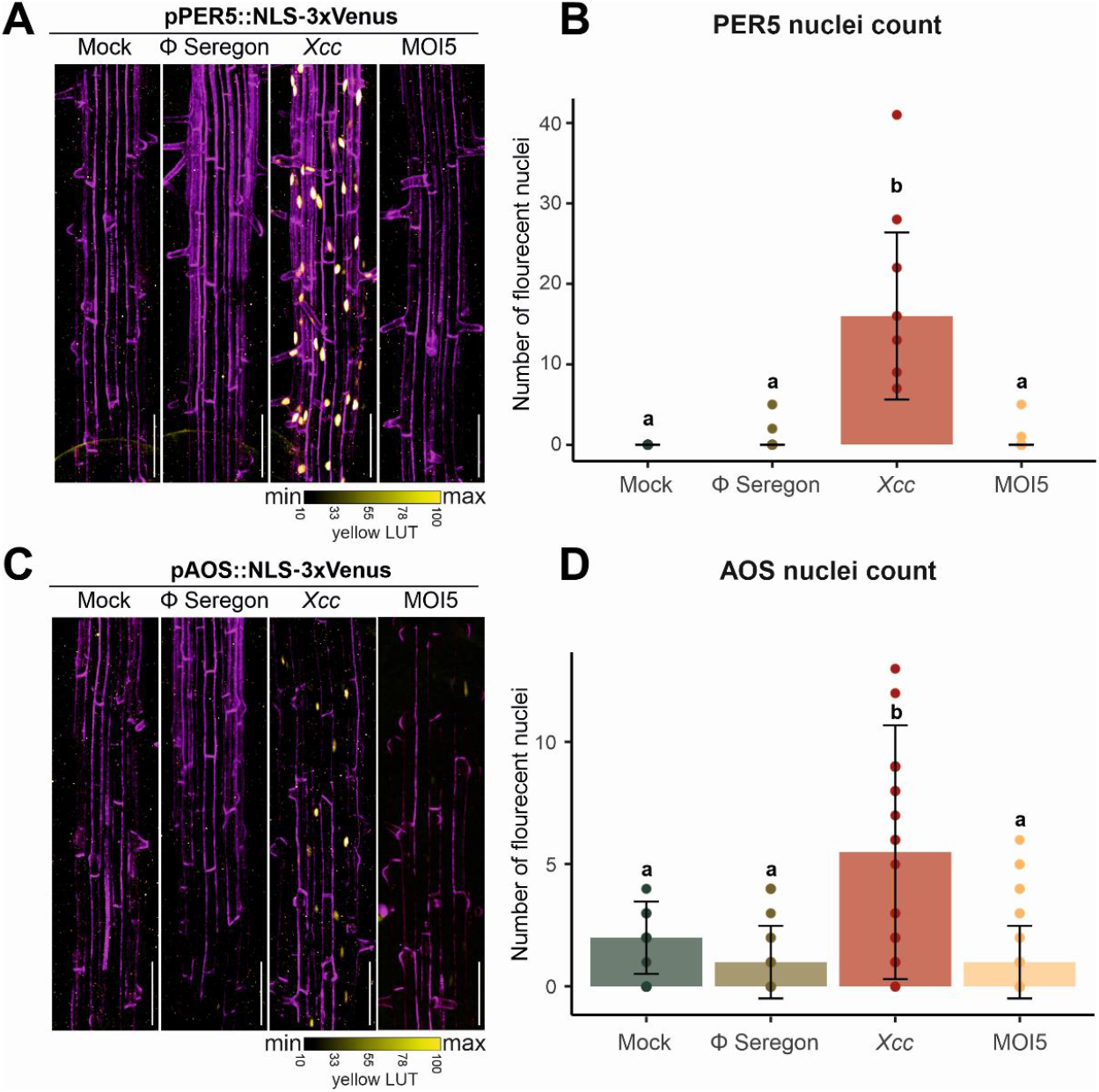
Nuclear localized reporter plant lines confirm reduced ROS and jasmonate response in phage-treated samples. **A)** Peroxidase superfamily reporter PER5 [69] induction (yellow) co-visualized with propidium iodide stained cell-walls (PI, magenta), in response to *Xcc*, phage Seregon, and MOI5 **B)** Nuclei count in PER5 reporter lines; Kruskal-Wallis chi-squared = 20.387, df = 3, p-value = 0.0001411. **C)** Jasmonic acid synthesis reporter line AOS (yellow) co-visualised with propidium iodide-stained cell-walls (PI, magenta response to *Xcc*, phage Seregon, and MOI5. Microscopic images are presented as maximum Z projection. **D)** Nuclei count in AOS reporter lines; Quantification of bright nuclei is given as a bar plot with median ±mad (median absolute deviation), differences were calculated Kruskal-Wallis chi-squared = 15.091, df = 3, p-value = 0.001741. Number of replicates is given on x-axis. Individual measurements are presented as data points. Scale bar= 100µm.

In similar manner, we tested if biocontrol will decrease JA synthesis gene expression as observed in the transcriptomics results. Here, the promoter of the gene encoding the JA synthesis protein AOS was coupled to the Venus reporter gene. Consistent with transcriptomic data, *Xcc*-treated roots showed strong JA-related signal accumulation, whereas phage biocontrol plants had significantly lower levels.

### 3.6 Phage biocontrol leads to reduced expression of bacterial virulence genes

To better understand how phage biocontrol influences *Xcc* during plant infection, we analyzed bacterial gene expression in the presence and absence of phage Seregon. PCA analysis revealed distinct bacterial transcriptomic profiles, with the strongest differences occurring at 2 dpi (Figure 5A). Differential expression analysis showed a significant shift in bacterial gene regulation under phage pressure (Figure 5B), with a marked reduction in virulence-associated genes and an increase in stress response pathways.

**Figure 5:**
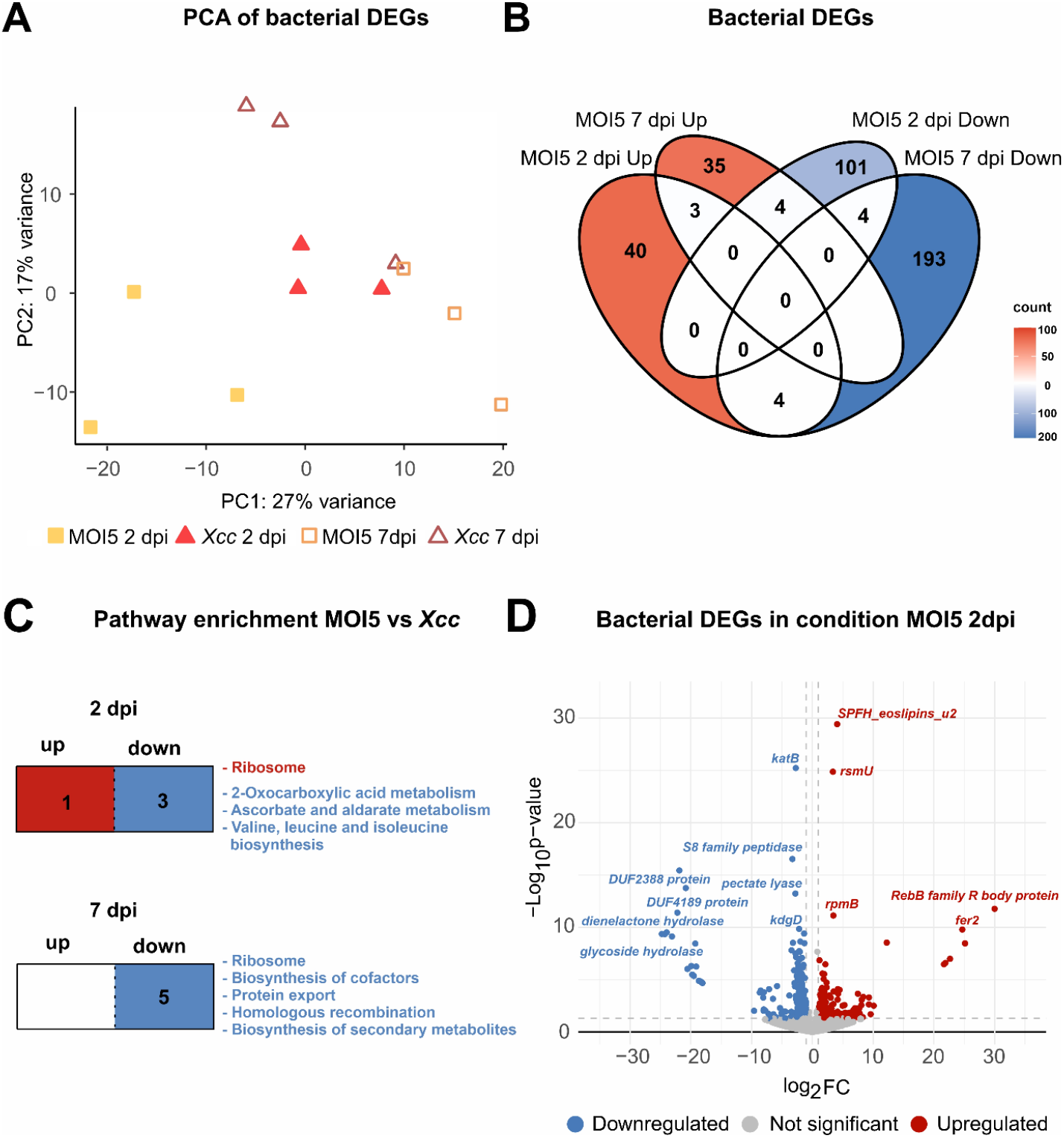
Phage biocontrol results in reduced expression of bacterial virulence-associated genes. Differentially expressed bacterial genes were identified with DESeq2; p-val < 0.05 **A)** Principal component analysis, samples: bacterial reads from samples MOI5 2 dpi, MOI5 7 dpi, *Xcc* 2 dpi and *Xcc* 7 dpi. **B)** VennDiagram of differential expressed genes in the presence of phages (MOI5), and the *Xcc*. **C)** Number of enriched pathways (Supplemental Table S4). **D)** Vulcano-plot of differentially expressed genes. Up-regulated genes are marked in red, down-regulated genes are marked in blue.

Remarkably, phage infection led to the downregulation of multiple *Xcc* virulence factors, particularly key T3SS components and effectors (*hrpF, xopX, xopE, xopAH*), likely hampering effector protein delivery required for plant colonization (Figure 6) [70]. Additionally, genes encoding enzymes involved in plant cell wall degradation (M91 metallopeptidase, pectate lyase, cellulase) showed reduced expression levels, along with *gspD*, a component of the T2SS secretion system, further limiting bacterial invasion. Reduced motility was indicated by the strong downregulation of *fliA*, the flagellar biosynthesis sigma factor, which may restrict systemic spread within *Arabidopsis* and is also associated with bacterial virulence. Interestingly, *Xcc* antioxidant defenses were also affected, with catalase genes *katB* and *katG* being significantly downregulated [71]. This may result from lower overall ROS levels in phage biocontrol plants. Another notable finding was the suppression of *qatB* and *qatC*, encoding components of the QatABCD anti-phage defense system. The observation that amongst the bacterial transcripts during biocontrol anti-phage systems were downregulated is counterintuitive at first glance, but might hint towards a phage-mediated suppression of bacterial defense [72].

**Figure 6.**
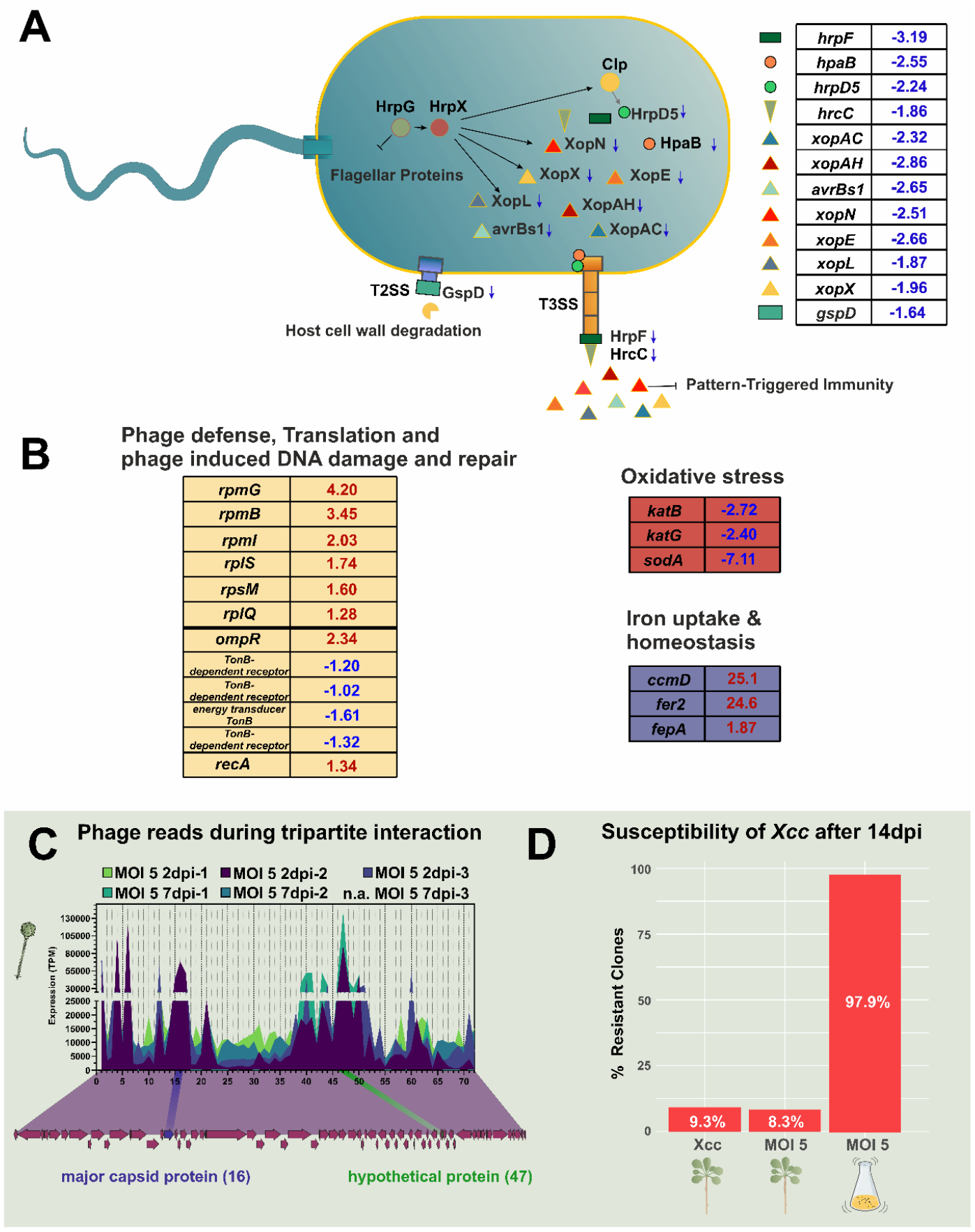
Bacterial virulence expression and phage transcripts. **A)** Significant differentially expressed effector genes (log2fold change) involved in *Xcc* virulence are presented in their regulatory context. **B)** Further functional bacterial clusters under *Xcc* versus MOI5 (log2fold change). **C)** Transcripts mapping to the genome of phage Seregon. **D)** Susceptibility testing to phage Seregon of *Xcc* colonies surviving after 14d in planta in presence and absence of phage compared to 14 d incubation in a batch in vitro culture.

### 3.7 Coexistence of phage Seregon and *Xcc* did not lead to enhanced emergence of bacterial resistance to phage infection

In our transcriptome analysis, we detected phage Seregon transcripts in infected plants (Figure 6C) at all time points. These results confirmed that phage biocontrol – as applied in our study – did not lead to the eradication of the pathogen *Xcc*, thereby enabling ongoing infection of *Xcc* by Seregon. Nevertheless, Seregon PFUs significantly declined over the course of the experiment (Figure 2E). Therefore, we wondered whether phage-treatment might lead to the emergence of bacterial resistance to phage infection. To test this, we isolated *Xcc* cells through an invasive harvest (Figure 2) and submitted surviving clones of *Xcc* to a phage sensitivity testing. Remarkably, only 6-10% of the surviving colonies retrieved from the plants were resistant to phage Seregon (Figure 6 D). In contrast, 97% of colonies isolated from an in vitro infection experiment were resistant. These results highlight the promising potential of phage-based biocontrol in reducing bacterial virulence while not leading to quick emergence of phage resistance in complex and structured environments like in the plant context. To identify the potential phage receptor, we sequenced four resistant *Xcc* clones resulting in 23 unique SNPs (Table S4). Mutations in a gene encoding a glycosyl transferase (XC_4244) and conserved hypothetical membrane protein DUF885 (XC_1704) provide first hints that glycosylation of cell wall components might be relevant for host recognition. Indeed, a knock-out of XC_4244 conferred partial resistance against phage Seregon (Figure S6).

## 4. Discussion

Bacteriophages remain an underutilized resource that bears a high potential for the development of sustainable biocontrol strategies against plant pathogenic bacteria. This potential arises from key characteristics of lytic phages: their host specificity and ability to self-propagate. In this study, we demonstrated that treatment with the lytic phage Seregon effectively protected *A. thaliana* seedlings from *Xcc* infection, restoring leaf area and plant weight to control levels. Our tripartite-transcriptomic analysis revealed that phage biocontrol reduced plant defense investment. While a single phage treatment did not completely eradicate *Xcc*, it significantly suppressed the expression of virulence genes in *Xanthomonas*, highlighting its potential as a biocontrol strategy.

Our study provides insights into the impact of phage biocontrol on plant immune signaling and bacterial adaptation at the level of gene expression. We found that phage biocontrol reduces bacterial effector secretion, allowing the plants PTI to effectively counter the pathogen. We observed alterations in ETI, hormonal signaling, and cell death regulation during phage biocontrol, which is in line with improved plant survival and performance under pathogen pressure. In agreement with these findings, phage-treated plants upregulate cytokinin biosynthesis (LOG8) and perception transcripts (AHP4, ARR3, ARR15) indicating a shift toward enhanced growth and development [73, 74].

Past studies already revealed the strong potential of bacteriophages bear in controlling crucifers’ black rot, from early proposals a century ago [4] to recent systematic testing of *Xanthomonas* phages in plant nurseries [75, 76]. Significant symptom reduction, effective seed decontamination, and lower bacterial load in treated plants have been identified [77]. Consistent with our findings, they also found that bacterial resistance emergence *in planta* remains limited [78–80]. Others observed reduced bacterial virulence in the phage resistant subpopulation of the pathogen [81, 82]. This highlights the importance of *in planta* studies, as *in vitro* approaches - albeit high-throughput - lack the physiochemical complexity of the plant environment. To improve future approaches, the design of complementary phage cocktails or phage engineering can further enhance biocontrol efficacy and mitigate resistance development [83].

The critical role of secretion systems for *Xanthomonas* virulence is well established [84–86]. In line with the restoration of plant growth, we observed significantly reduced expression of virulence-associated genes in phage-treated samples. Specifically, we detected downregulation of the T2SS outer membrane translocon GspD, which is important for transporting folded proteins into the periplasm [87]. In addition, genes encoding components of the T3SS translocon HrpF and several T3SS effectors showed reduced expression. Among these, the secreted effector XopN - previously shown to interfere with receptor-like kinase (RLK) function in tomato and thereby hampering PTI [88] – was notably reduced, further supporting the impact of phage biocontrol in weakening bacterial virulence at the plant environment.

While most studies have focused on the interactions between bacteria and phages or between bacteria and plants, the potential for a direct interaction between plants and phages remains unknown. In mammalian models, phages exhibit niche specialization and influence immune modulation [89, 90]. Studies have shown that phages adhere to the human mucus, enabling their persistence within this niche [90, 91]. Likely similar and highly complex niche adaptations occur for phages in the plant environment [7]. Interaction with “non-host” organisms, like fungal hyphae, was further shown to be involved in the transport of phages to new soil habitats [92]. First attempts also studied phage translocation of phages in the plant system, but did not yet provide mechanistic insights [93, 94]. Notably, the coexistence of bacteria and phages was not driven by a widespread development of phage resistance within the bacterial population in planta. This suggests that phage resistance entails fitness trade-offs for the pathogen in the host environment. It remains an intriguing question whether plants have evolved mechanisms that modulate phage activity, as recently proposed for bacterial communities [95].

Altogether, our study provides comprehensive transcriptional insight into phage-based biocontrol of bacterial plant pathogens. While our parallel transcriptomics approach represents a valuable first step, future research should further elucidate the molecular mechanisms underlying viral perception and immune homeostasis within both synthetic and native plant-microbe consortia. A deeper molecular understanding of these complex interactions is essential for the rational design of effective and sustainable phage-based biocontrol strategies.

## Contributions

S.E., B.A., U.S. and J.F. designed the study. S.E. and M.Z. performed the experiments. S.E. and B.A. performed the RNA-Seq data analysis. M.Z. performed microscopy and image analysis. S.E., G.G., J.F., and B.A. wrote the manuscript. B.A., U.S., G.G. and J.F. supervised the experiments. All authors read, revised, and approved the final manuscript.

## Acknowledgments

The authors gratefully thank Prof. Dr. Franz Narberhaus (RUB, Bochum) for providing the *Xanthomonas campestris* 8004 strain, Michaela Gerads for technical assistance and all teams for the supportive working environment.

