## Supplementary Material for "Phage biocontrol reduces the burden on plant immunity through suppression of bacterial virulence"

Summary:

The Supplementary information includes 5 figures and 7 Tables

Figures

**Figure S1: *Arabidopsis* defence regulation in different microbial treatments compared to control plants**

**Figure S2. Phage biocontrol (MOI5) leads to altered defence regulation in *Arabidopsis* compared to *Xcc* infected plants.**

**Figure S3: Phage biocontrol leads to reduced expression of virulence-associated genes in *Xanthomonas campestris (Xcc).***

**Figure S4: Bacterial survivors after at the end of the in-planta experiment and phage susceptibility testing**

**Figure S5: Mutation of XC_4244 encoding a glucose-6-phosphate 1-epimerase significantly reduces susceptibility of *Xcc* to infection with phage Seregon.**

Tables

| | |
|---|---|
| **Supplemental Table 1** | **KEGG Pathway enrichment Arabidopsis.xlsx** |
| **Supplemental Table 2** | **Arabidopsis AGI codes and ratios-Figure 3 and Figure S1.xlsx** |
| **Supplemental Table 3** | **Xanthomonas DEGs in functional context.xlsx** |
| **Supplemental Table 4** | **KEGG Pathway enrichment Xanthomonas.xlsx** |
| **Supplemental Table 5** | **Xanthomonas SNPs.xlsx** |
| **Supplemental Table 6** | **Raw DEGs.xlsx** |
| **Supplemental Table 7** | **Strains used in this study** |
| **Supplemental Table 8** | **Primers used in this study** |

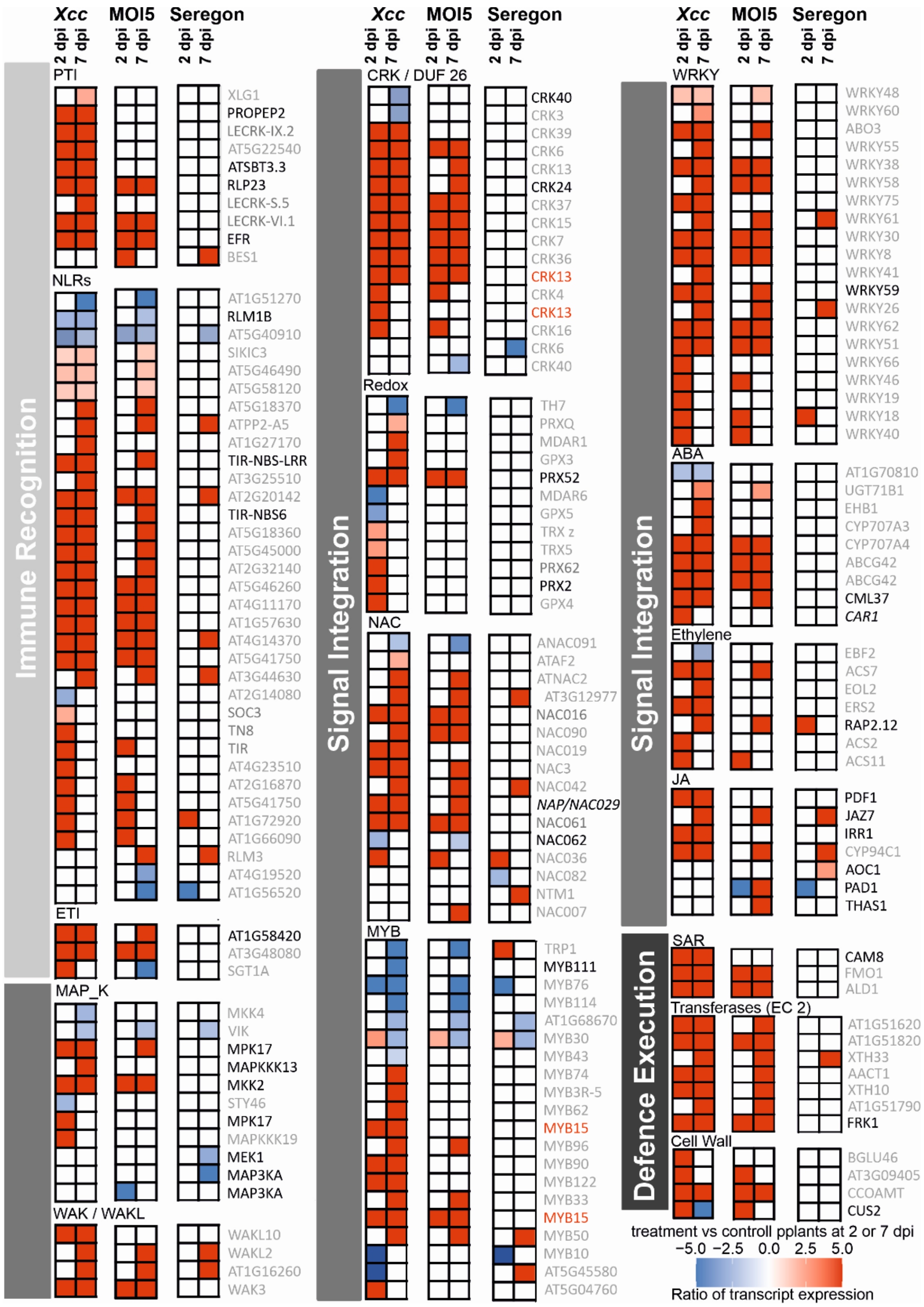


**Figure S1: *Arabidopsis* defence regulation in different microbial treatments compared to control plants**

Ratio of gene expression of the 3 treatments (Xcc, MOI5 and Seregon) compared to control plants (expressed as log2 (FC)). Shown are selected transcripts thought to be involved in pathogen recognition (light grey), immune signal integration (medium grey) and defense execution (dark grey). The level of gene expression is presented as a heat map, shown bottom right, with shades of blue indicating lower expression, and shares of red indicating higher expression. The heat map is cut off at +/-5 for visual simplification, numerical data can be found in supplemental table S2. Transcripts are grouped in functional groups based on MapMan mapping file X4.1 [1], with adaptations as listed in supplemental table 2. Transcript names mentioned in the text are in black, and additional names are in grey, names of splice isoforms of the same transcript are in red.

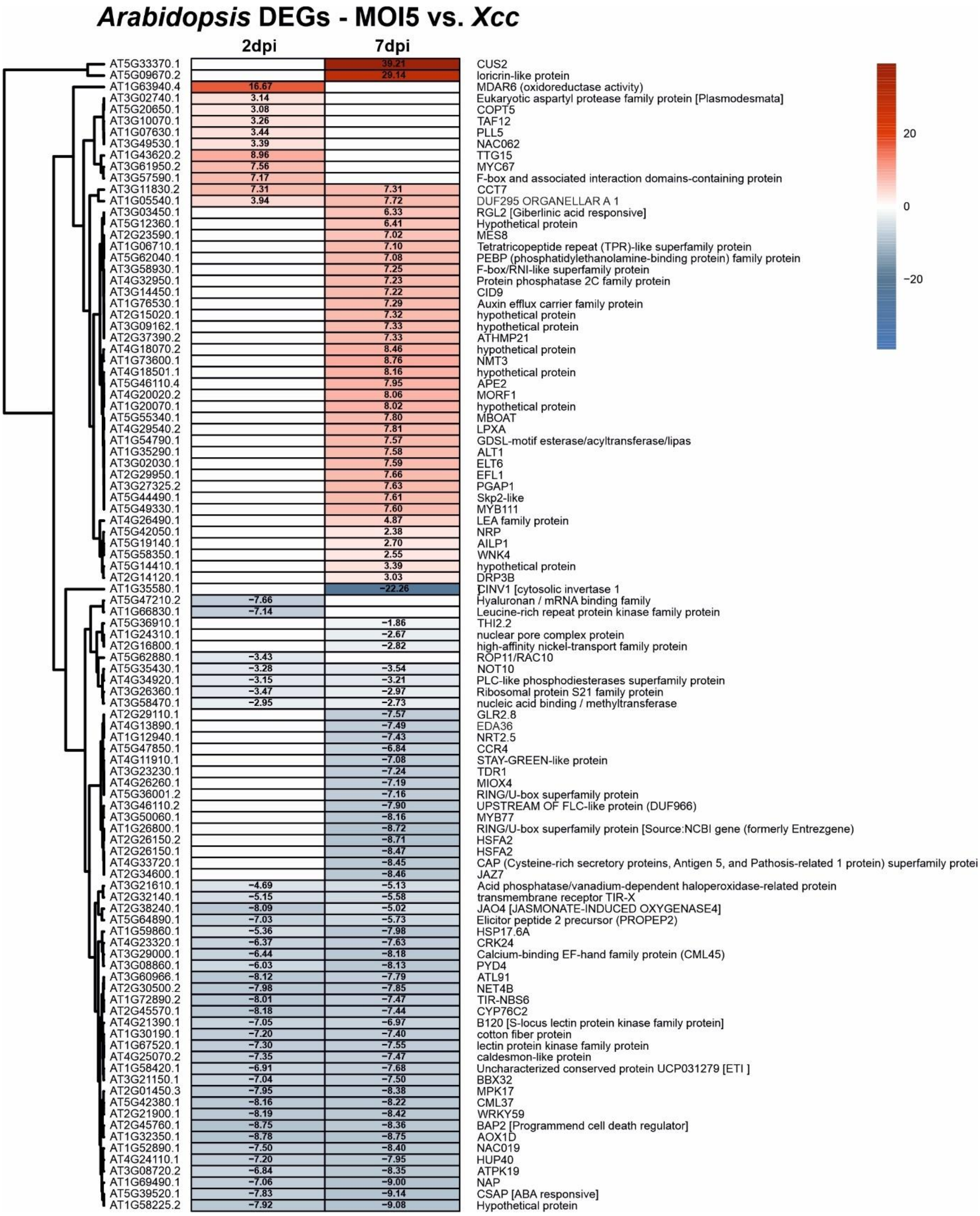


**Figure S2. Phage biocontrol (MOI5) leads to altered defence regulation in *Arabidopsis* compared to *Xcc* infected plants.** Global *A. thaliana* transcriptome analysis during the tripartite interaction. Heat map of the 100 most significant differentially expressed plant genes. Gene names based on TAIR entries were added. Shown are log2fold changes for three independent biological replicates MOI5 vs *Xcc* at 2 or 7 dpi. Empty cells are not differentially expressed between the two conditions (MOI5 versus *Xcc*) at the given day.

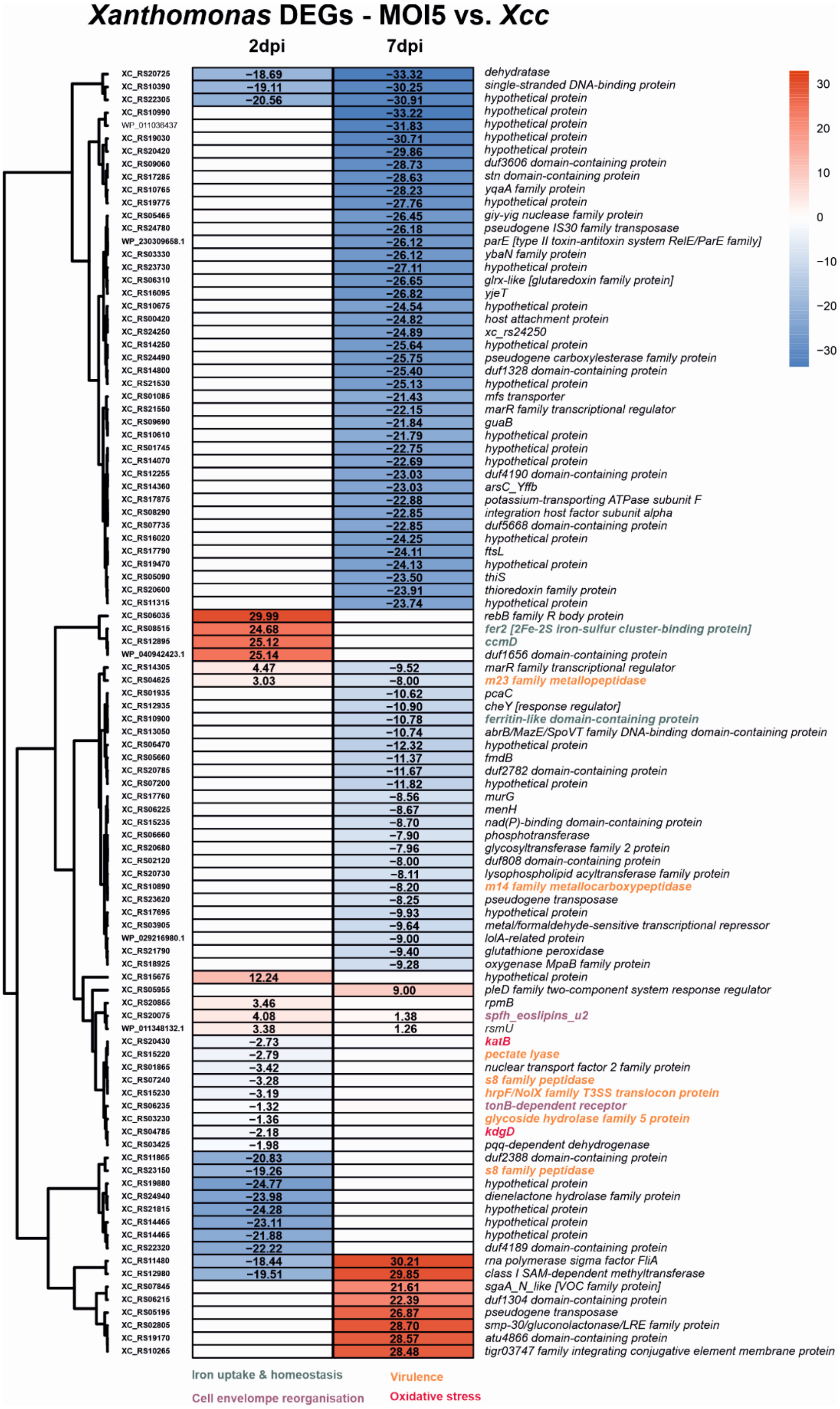


**Figure S3: Phage biocontrol leads to reduced expression of virulence-associated genes in *Xanthomonas campestris (Xcc).***

Global bacterial transcriptome analysis during the tripartite interaction. Heat map of the 100 most significant differential expressed bacterial genes in presence or absence of the phage. Gene identifiers were replaced with gene names based on NCBI entries. Shown are log2fold changes of data for three independent biological replicates (MOI5 versus *Xcc* at 2 or 7 dpi).

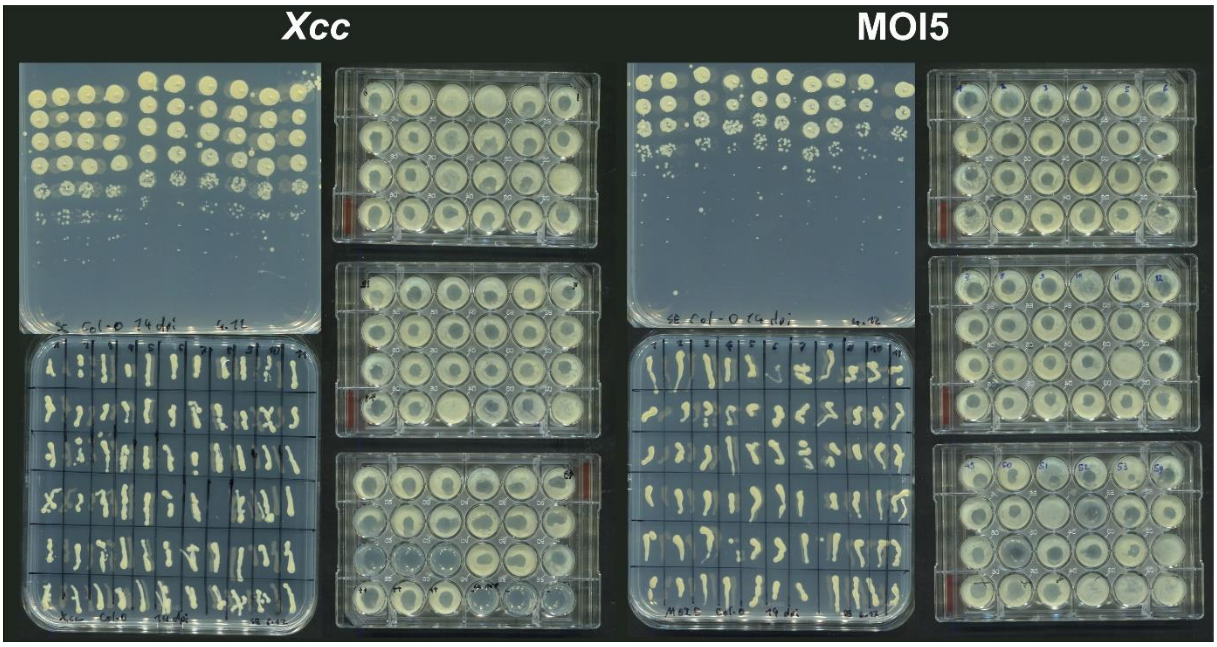


**Figure S4: Bacterial survivors after at the end of the in-planta experiment and phage susceptibility testing**
After 14 days of growth in the plant environment (on ½ MS plates) individual colonies extracted from plant material. Those “survivors” were re-streaked on nutrient agar and subjected to a 48 well double agar overlay spot assay. Susceptible clones show lysis zones whereas resistant ones don’t.

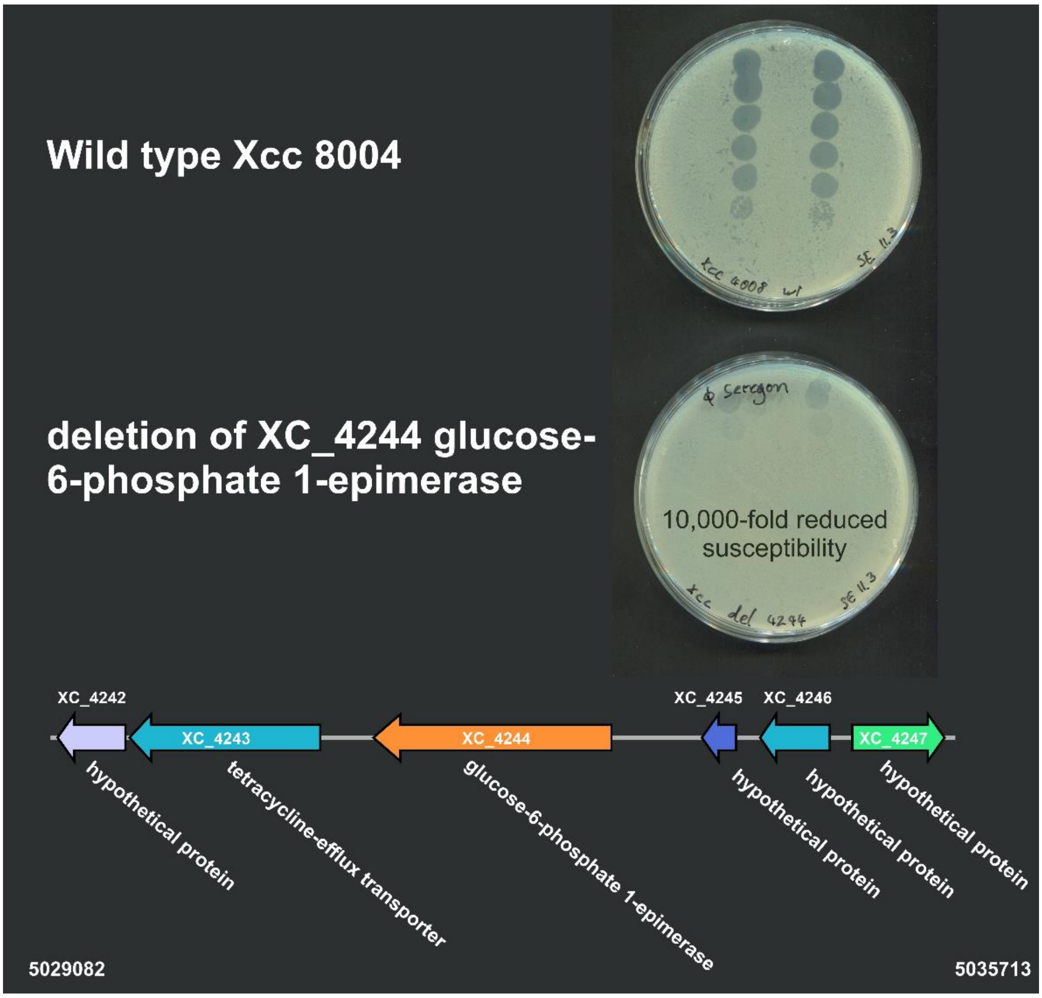


**Figure S5. Mutation of XC_4244 encoding a glucose-6-phosphate 1-epimerase significantly reduces susceptibility of *Xcc* to infection with phage Seregon.**

To test if the SNPs identified in "bacterial survivors" of the phage biocontrol treatment were relefant for phage infection, we constructed a knockout mutant of XC_4244 by homologous recombination. *Xcc* 8004 wt and *Xcc*ΔXC_4244 were infused into the 0.4%-soft agar of a double agar overlay. Dilution series of *Xanthomonas* phage Seregon was spotted on top and incubated for 24 h at 28°. Displayed below is the genomic context of gene XC_4422.

**Supplemental Table 7 Strains used in this study**

| Organism | Source | Reference |
|---|---|---|
| *Xanthomonas campestris* pv. *Campestris* 8004 | AG Narberhaus (RUB, Bochum) | [2] |
| *Xanthomonas campestris* pv. *Campestris* Δ XC_4244 | This study | - |
| *Xanthomonas* phage Seregon | Erdrich et al. 2022 | [3] |

**Supplemental Table 8 Primers and plasmids used in this study**

| Primer Name | Sequence | Reference |
|---|---|---|
| InFusion_Xcc_4244_del_LF_F | cgccaagcttgcatgcctgcagatcaggagagatggacat | This study |
| InFusion_Xcc_4244_del_LF_R | ccttaattctctagttgagaccccaagacagtaaagaggca | This study |
| InFusion_Xcc_4244_del_RF_F | ataaagtattgagaccaatcaggcaggcgtct | This study |
| InFusion_Xcc_4244_del_RF_R | gtaaaacgacggccagttaacacaccgacgtca | This study |
| InFusion_Xcc_4244_Kan_F | ggtctcaactagagaattaagggag | This study |
| InFusion_Xcc_4244_Kan_R | ggtctcaatactttatcctagtttg | This study |
| Seregon_Capsid-F | atggatttgttcagcactgcgg | This study |
| Seregon_Capsid-R | agcttcagctcgtcgaacatg | This study |
| pK19mobsacB | | [4] |
